# Quantitative fluorescent nanoparticle tracking analysis and nano-flow cytometry enable advanced characterization of single extracellular vesicles

**DOI:** 10.1101/2024.04.30.591813

**Authors:** Danilo Mladenović, Joseph Brealey, Ben Peacock, Nataša Zarovni

## Abstract

Current state-of-the-art tools for analyzing extracellular vesicles (EVs) offer either highly sensitive but unidimensional bulk measurements of EV components, or high-resolution multiparametric single particle analyses which lack standardization and appropriate reference materials. This limits the accuracy of assessment of marker abundance and overall marker distribution among individual EVs, and finally, the understanding of true EV heterogeneity.

In this study, we aimed to define the standardized operating procedures and reference material for fluorescent characterization of EVs with two commonly used EV analytical platforms - nanoparticle tracking analysis (NTA) and nano-flow cytometry (nFCM).

For the first time, we achieved quantitative fluorescence analyses on ZetaView NTA and NanoAnalyzer nFCM instruments, by utilizing yellow-green FluoSpheres (FS) with assigned ERF (equivalent reference fluorophore) values. This standardization technique allowed for fluorescent EV signal to be expressed in ERF units (indicative of bound fluorescent antibodies per EV), thus enabling measurement of target protein marker abundance on individual EVs, and in the whole EV population. The NTA’s and nFCM’s limits of quantification (LoQ) were evaluated at 115 and 75 Alexa Fluor 488 (AF488) molecules, respectively. To complement these shortcomings, in-line bulk fluorescence measurements in a plate reader were performed. This provided absolute marker quantification, and more insightful analyses of EV heterogeneity and marker stoichiometry.

The standardization method outlined in this work unlocks the full analytical potential of NTA and nFCM, enabling cross-platform data comparison. At the same time, it highlights some of the technical challenges and considerations, and thus contributes to the ongoing efforts towards development of EV analytical tools.

## INTRODUCTION

Over the past decade, we have witnessed a rise, and a great expansion of advanced platforms and tools for analyses of extracellular vesicles (EVs) [1]. From bulk assays, which usually provide high sensitivity but unidimensional measurements, the analyses have reached down to a single particle level, opening the possibility for multiparametric assessment of individual EVs, and deeper understanding of their heterogeneity and complexity [2–5]. These systems allow simultaneous biophysical and biochemical evaluation of EVs by measuring their size, concentration, charge, morphology, refractive index, and/or composition. The latter is often achieved via fluorescent reagents that target different components (lipids, proteins, nucleic acids, metabolites, etc.), present either on the membrane or within the lumen of EVs [6,7]. A common strategy, offering great flexibility and value for EV characterization, utilizes antibodies labeled with various fluorochromes, to detect a rich display of proteins on the EV surface. This enables phenotyping of different EV subpopulations, to determine their putative cellular or subcellular origin, and also offers the potential for biomarker discovery, making possible EV-based diagnostics [8,9].

Even though state-of-the-art instruments for single particle analyses offer higher resolution compared to the bulk assays, their true potential for EV phenotyping is often downplayed by the lack of standardized protocols and reference materials. Popular methods, such as NTA, nFCM, or even super-resolution microscopy and on-chip fluorescence imaging capable of single-molecule sensitivity, are currently achieving qualitative or semi-quantitative analyses, at best [6,10–13]. Quantitative single molecule localization microscopy (qSMLM) possesses unparalleled analytical power for quantification of marker abundance per individual EVs [14,15]. However, fluorescent NTA (FNTA) and nFCM offer greater flexibility and high throughput multiparametric analyses, making them preferred tools for day-to-day use in EV research. Even if more conventional, FNTA and nFCM are still not standardized for fluorescent quantification of EV content. Choice of calibrators, which would unlock their full quantitative potential, is reduced to the well-established MESF (molecules of equivalent soluble fluorophore) beads, used in the traditional flow cytometry [16,17]. However, the fluorescence they emit is orders-of-magnitude brighter than the fluorescent signal we might expect from any labeled EV. In turn, this requires data extrapolation outside of the calibrated quantification range, producing estimates which have much higher uncertainty [16,18]. Smaller and dimmer hard-dyed beads assigned with ERF values may provide a suitable alternative for fluorescence calibration. However, they usually have spectrally mismatched multi-peak emission, with respect to the fluorochromes used in EV labeling, which limits their application [19].

In this work, we defined the analytical protocols and evaluated the performance of two commonly used instruments for single EV analyses – Particle Metrix ZetaView and NanoFCM NanoAnalyzer. We employed yellow-green FS hard-dyed nanoparticles, with single-peak FITC-like emission spectra, for standardizing fluorescence readout in ERF units and estimating the instruments’ LoQ. To demonstrate the method’s utility, EVs purified from simple (cell conditioned media) and complex (blood plasma) biofluids were used for analyzing multiple markers and their abundance per single EV and in the whole EV population, via AF488-labeled antibodies. Finally, we complemented single particle analyses of FNTA and nFCM with high-sensitivity bulk fluorescence measurements performed in a plate reader, to surpass each platform’s individual limitations and obtain more comprehensive results.

## MATERIALS AND METHODS

### Extracellular vesicles derived from COLO cell line

Colorectal cancer cell line-derived small EVs, referred to as COLO sEVs (Lyophilized Exosomes from COLO1 cell line; HansaBioMed Life Sciences, Estonia), and medium size EVs, referred to as COLO mEVs (Lyophilized Microvesicles from COLO1 cell line; HansaBioMed Life Sciences), were used for staining with fluorescent antibodies.

### Blood plasma processing and EV purification

Healthy donor blood plasma pool (50 donors), containing EDTA as an anticoagulant, was purchased from BioIVT (Human Plasma K2EDTA; BioIVT, United Kingdom). The plasma was obtained from the whole blood within 15 minutes after the collection, by centrifugation at 2,800 ×g for 20 minutes, in a refrigerated centrifuge (5°C). Afterwards, it was frozen and shipped with the dry ice. Prior to the experiments, the plasma was thawed and immediately centrifuged for 15 minutes at 2,500 ×g, at 4°C, followed by filtration through a 0.45 µm PES syringe filter (Minisart; Sartorius, Germany). Such precleared plasma was then used for EV purification.

Size exclusion chromatography column PURE-EVs (HansaBioMed Life Sciences) was equilibrated with PBS. For EV purification, 2 mL of precleared plasma were loaded on top of the column, and fractions of 0.5 mL were collected. Fractions 6-11 containing EVs were pooled and used for staining with fluorescent antibodies.

### EGFP fluorescent EVs

Commercially available EVs derived from HEK293 cell line overexpressing CD63-EGFP (FLuoEVs; HansaBioMed Life Sciences) were used for direct analyses on FNTA to define the traces-to-particles (T:P) ratio criteria. Furthermore, FLuoEVs were used to assess the effect of different FNTA threshold levels (minimum brightness) on event detectability and overall analysis outcome.

### Silica and polystyrene beads

Monodisperse 105.1 nm (Corpuscular, Canada) and polydisperse 68, 91, 113, and 155 nm (1:1:1:1 ratio, NanoFCM Inc., China) size standard silica beads were used for comparison of particle concentration and size measurements in scatter mode of NTA and nFCM.

Yellow-green FluoSpheres carboxylate-modified microspheres of 100, 40, and 20 nm (Thermo Fisher Scientific, United States) were used for antibody assignment. Subsequently, they were used as a reference material for fluorescence analyses on FNTA, nFCM, and plate reader. Detailed description of methodology is provided in the separate sections.

### Staining EVs with fluorescent antibodies

EVs were stained using the antibodies described in the Table 1.

**Table 1.** Antibodies used for EV staining.

| Sample | Antibody description | Provider |
| --- | --- | --- |
| COLO sEVs and<br>COLO mEVs | Human CD9 Alexa Fluor 488-conjugated antibody, IgG2B | R&D Systems |
|  | Human CD63 Alexa Fluor 488-conjugated antibody, IgG1 |  |
| Plasma EVs | Human CD9 Alexa Fluor 488-conjugated antibody, IgG1 | Exbio |
|  | Human CD63 Alexa Fluor 488-conjugated antibody, IgG1 |  |
|  | Human Integrin alpha 2b/CD41 Alexa Fluor 488-conjugated antibody, IgG1 | R&D Systems |
|  | Human Glycophorin A Alexa Fluor 488-conjugated antibody, IgG1 |  |

Staining reactions for fluorescence analyses were prepared according to the Table 2.

**Table 2.** EV staining reactions.

| Staining reaction | Volume [μL] | Number of particles | Antibody [μg] |
| --- | --- | --- | --- |
| COLO sEVs | 40 | $5 \times 10^9$ | 0.8 |
| COLO mEVs | 40 | $2.5 \times 10^9$ | 0.8 |
| Plasma EVs | 100 | $5 \times 10^{10}$ | 0.25 |

Incubation was carried out at 37°C (for COLO EVs) or at room temperature (for plasma EVs), covered from light, under continuous low-speed shaking for 1.5 hours. Afterwards, the unbound antibodies were removed as previously described [6]. Briefly, samples were washed with 450 µL PBS using Nanosep centrifugal devices with Omega membrane 300K (Cytiva, United States) and centrifugation at 3,000-5,000 ×g, until most of the liquid had passed in the permeate. After 6 washing cycles, the retentate was recovered in ∼100-110 µL of PBS and analyzed on NTA, nFCM, and plate reader. PBS only, dye only, and EVs only (unstained) were used as process controls and for background subtraction.

### Assignment of antibody/AF488/EGFP equivalents to FS beads

Yellow-green FS beads and AF488-labeled antibodies were serially diluted in PBS and loaded in a black polystyrene 96-well plate with no binding capacity (Biomat, Italy). The dilutions were adjusted in a way that the measured fluorescence intensity of the FS beads fits in the range of the fluorescence intensity of the serially diluted antibodies. The measurements were performed on a GENios Pro microplate reader (Tecan Group Ltd., Switzerland) with 485 nm excitation wavelength, and emission was detected through 535/25 nm bandpass filter. The number of antibodies per well was calculated based on the known antibody concentration (µg/mL), molecular weight (IgG ∼150 kDa = 150,000 g/mol), and Avogadro’s constant (6.022×10^23^ molecules/mol). After subtracting the blank (PBS), the plate reader data (fluorescence intensity), the number of beads per well, and the number of antibodies per well were log-transformed, and linear regression was plotted using antibodies as a reference. From the obtained equation, the fluorescence intensity of the FS beads was converted to the number of equivalent fluorescent antibodies. This was done for each individual antibody type and lot, due to the differences in antibody production and subsequent fluorophore-to-protein (F:P) ratios, i.e., different amount of conjugated AF488 molecules per antibody. All of the serial dilutions and measurements were performed in at least three independent experiments, to obtain the average number of antibody molecules per bead (Supplementary file 1). The standard deviation was on average 9% (2-15% range). Following the principles previously reported [20–22], additional correction factors were applied to these calculations to compensate for the differences in optical configurations between the instruments (plate reader, FNTA, and nFCM), and slight spectral mismatching of AF488 and FS beads. The excitation/emission spectra of fluorophores, and excitation and collection efficiencies of different optical configurations, were obtained from https://www.fpbase.org/spectra/ [23,24]. For FNTA, correction factor of ×1.17 was applied to the number of antibodies assigned per each FS bead using plate reader, while for nFCM, correction factor was ×1.04. See the Supplementary files 1 and 2, and Figure S1 for more information.

Using known F:P ratio for multiple different antibodies (provided by the antibody manufacturers, Supplementary file 1), the assigned antibody equivalents per each FS bead were converted to AF488 equivalents.

To further convert the signal to EGFP equivalents per FS bead, the intensity and spectral differences between AF488 and EGFP were taken into account, and the correction factor of ×1.66 was calculated for FNTA optical configuration (Supplementary files 1 and 2).

### Nanoparticle tracking analyses

Particle Metrix Zeta View PMX-120 NTA instrument with 488 nm laser (Particle Metrix, Germany) was used for single-particle analyses in scatter and fluorescence mode. Instrument setup was done according to the manufacturer’s guidelines (cell check and auto-alignment with 100 nm polystyrene beads standards upon each start-up). For analyses in scatter mode, camera sensitivity was set at 85, shutter speed at 100, with high video quality at 30 frames/second, minimal area 10, maximal area 10,000, minimum brightness 20, and minimum tracelength 15. Fluorescence measurements of FS beads and fluorescent EVs were carried out using 500 nm long-pass filter, with the camera sensitivity set at 95, shutter speed at 100, with low video quality at 30 frames/second, minimal area 10, maximal area 1,000, minimum brightness varying between 20, 25, 30, and 35 (depending on the background level, and desired T:P ratio), minimum tracelength set at 5, and with “Low Bleach” technology enabled. PBS was used as a diluent for every NTA analyses.

FS beads were measured in each experiment using the same protocol as for stained EVs (described above), and FCS files were exported. In the case of multiple measurements of a single sample, FCS files were merged using an online tool floreada.io/fcsmerge. The FCS files were then opened using floreada.io/analysis, and custom parameter for fluorescence intensity ‘FI[log10]’ was created using the formula *log10({Mean Intensity}*{Area})*. The gating was performed for each bead population. This was particularly necessary for FS beads of 100 nm since their fluorescence intensity was too high, causing the beads that are out of focus to still “peek” into the focal plane, appearing as dimmer events (Figure S2). Median fluorescence intensity (MFI) was calculated for each gated population and statistics were exported (Save CSV statistics file). MFI was then used together with the assigned antibody, AF488, or EGFP values corrected for the FNTA (see Assignment of antibody/AF488/EGFP equivalents to FS beads), and linear regression was plotted. The equation thus obtained was then used to estimate the number of bound antibodies (type and lot specific), AF488, or EGFP molecules, respectively, per each traced fluorescent event. The example of excel formulae used for analyses of FCS files of stained EV samples is provided in the Supplementary file 3.

When analyzing stained EV samples in fluorescence mode, minimal criteria were established – to acquire at least 300 total traces per analyzed sample, and the T:P ratio had to be ≥35%, comparable to the one obtained in scatter mode. The former was achieved by running multiple acquisitions and injecting more concentrated sample into the FNTA cell (if the fluorescent events were scarce, like in plasma EV samples).

The desired T:P ratio was obtained by reanalyzing the video file of a same sample with a minimum brightness set at 20, 25, 30, or 35. Finally, the traces collected in FCS files were used for further analyses.

For the results summary of scatter measurements, particle concentration (subtracted by blank) was obtained from the 11-position table, while median and average particle size were calculated from the FCS files. For fluorescence measurements, particle concentration was subtracted by blank and additionally unstained samples, if there were any detectable autofluorescent particles. The median and average particle size, and the number of antibodies per particle were computed from the FCS files using the excel formulae in the Supplementary file 3.

### Nano-flow cytometry

A NanoAnalyzer U30 instrument (NanoFCM Inc.) equipped with dual 488/640 nm lasers was used for simultaneous detection of side scatter (SSC) and fluorescence of individual particles. Single-photon counting avalanche photodiode detectors (SPCM APDs) with bandpass filters allowed for collection of light in specific channels (SSC - 488/10 nm; FL1 – 525/40 nm; FL2 – 670/30 nm). The sampling pressure by air pump module was 1 kPa, sheath fluid was gravity fed HPLC-grade water, and measurements were taken over 60 seconds. Values for peak height (mean + 3 standard deviations of the background) and peak width (0.3 ms) were used as thresholds for peak identification.

For each particle, peak area was recorded in all three detection channels simultaneously, for use in constructing dot plots and histograms. Samples were diluted to attain a particle count within the optimal range of 1,500-12,000/minute. Blank measurements of TE buffer (10 mM Tris, 1 mM EDTA) were recorded, containing 200-400 particles/minute; these particles were subtracted from the sample measurement for concentration calculation. In the fluorescence measurements, additional subtractions were performed using the unstained control samples, if there were any detectable autofluorescent events.

Particle concentrations were determined by comparison to a standard containing 250 nm silica nanoparticles of known concentration. Particles were sized according to standard operating procedures using a proprietary 4-modal silica nanosphere cocktail (NanoFCM Inc., S16M-Exo). Using the NanoFCM software (NanoFCM NF Profession V2.0), a standard curve was generated based on the intensity of side scattered light of the four different silica particle populations of 68, 91, 113, and 155 nm in diameter. The laser was set to 15 mW and 10% SSC decay.

FS beads were measured in each experiment using the same protocol as for the stained EVs. MFI of each bead population was used in combination with the assigned antibody values corrected for the nFCM (correction factor ×1.04; see Assignment of antibody/AF488/EGFP equivalents to FS beads), and linear regression was plotted. The equation thus obtained was used to estimate the number of bound antibodies (type and lot specific) per each detected fluorescent event.

Data processing was handled within the NanoFCM NF Profession V2.0 software, with dot plots, histograms, and statistical data provided in a single PDF. Gating within the software allows for proportional analysis of subpopulations separated by fluorescence intensities.

### Microplate reader

The bulk fluorescence measurements were performed using GENios Pro microplate reader (Tecan Group Ltd., Switzerland) with 485 nm excitation wavelength, and emission was detected through 535/25 nm bandpass filter. Stained and washed EVs (5×10^8^-8×10^9^ particles/well, depending on the sample) were loaded in a black polystyrene 96-well plate with no binding capacity (Biomat, Italy) and diluted with PBS up to 100 µL. Same number of unstained EVs or the same volume of blank controls were loaded in the plate and used to subtract the background. All controls produced a negligible amount of background noise, i.e., comparable to PBS. As a reference, serially diluted FS beads were loaded and measured in parallel to produce standard curve (with the log-transformed data) based on the previous antibody assignments. The equation of this linear regression was then used to calculate the number of antibodies in our samples (expressed as antibodies/µL of sample). For comparison, antibody concentration in a sample was also estimated from FNTA and nFCM measurements by multiplying the average number of antibodies/particle (average epitope abundance) with the fluorescent particle concentration.

### Data analyses

FCS files from FNTA were analyzed using the Floreada.io online tool and excel formulae in supplementary file 3. NFA files from nFCM were analyzed using the NanoFCM NF Profession V2.0 software. All the results are shown as the mean ±SD, n≥3. The data were analyzed and the figures were plotted using GraphPad Prism 10.2.1.

## RESULTS

### Comparative detection of synthetic nanoparticles across EV analysis platforms

To first compare the performance of NTA in scatter mode (SNTA), and nFCM in particle sizing and quantification, monodisperse (105.1 nm) and polydisperse (68, 91, 113, and 155 nm in a 1:1:1:1 ratio) size standard silica beads were analyzed. For this purpose, the same settings were used as for the analysis of EVs in scatter mode. Both instruments produced similar measurements of concentration and size for monodisperse beads, although the latter was significantly lower compared to the beads rated value (Figure 1A). This discrepancy can be explained by the fact that the beads’ rated size comes from a DLS measurement, which is a method known to provide a bias towards larger particles [25]. With polydisperse silica beads, the concentration appeared significantly higher on SNTA (Figure 1B). This observation is in line with prior reported overestimation in SNTA concentration measurements of polydisperse samples [26]. Nevertheless, the estimated median size remained similar between SNTA and nFCM, and concordant with the rated value.

**Figure 1.**
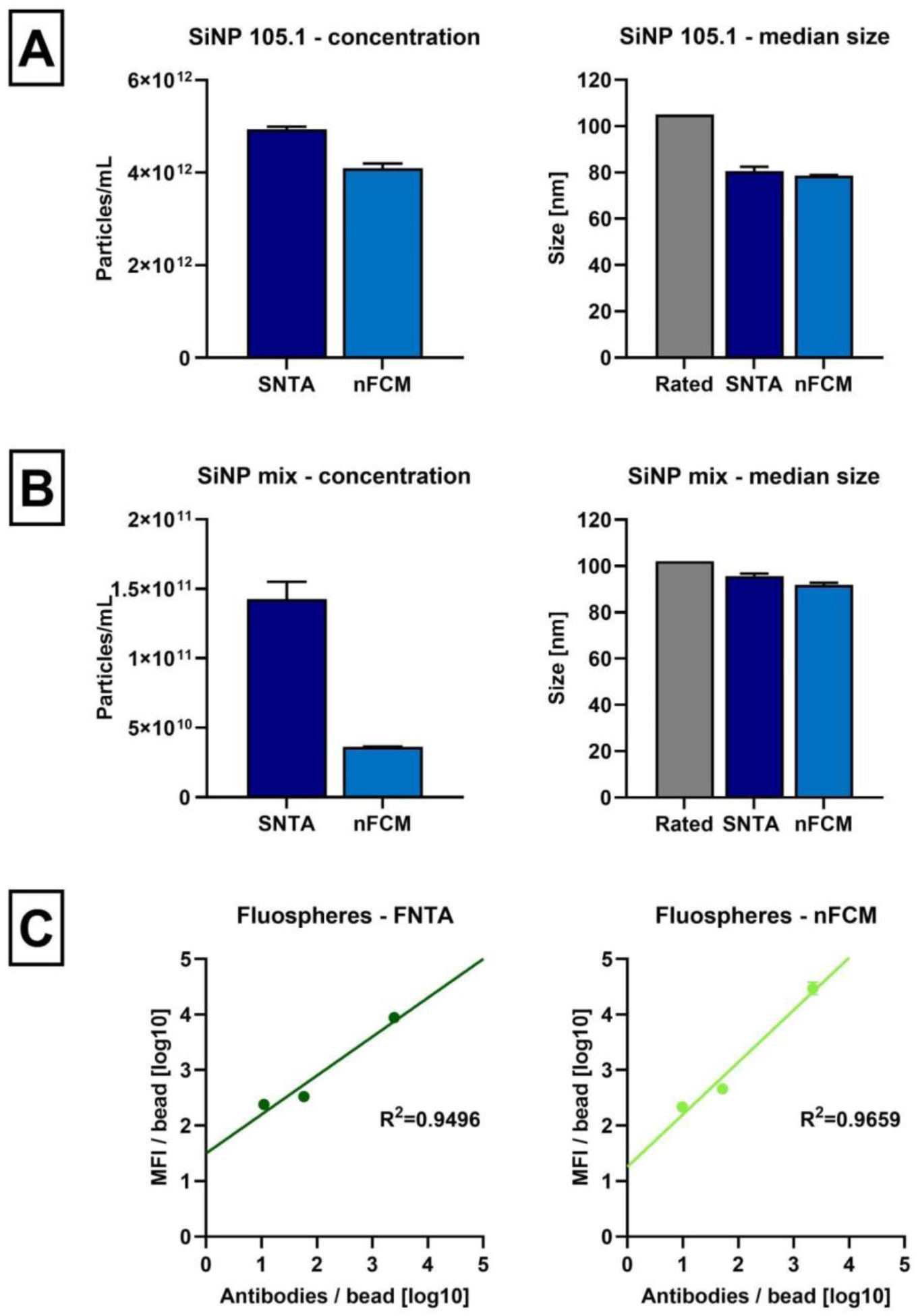
Silica and FS beads. Silica bead size standards in **(A)** monodisperse (105.1 nm) and **(B)** polydisperse suspensions (68, 91, 113, and 155 nm in a 1:1:1:1 ratio) were measured on SNTA and nFCM. **(C)** FS beads of different sizes (20, 40, and 100 nm) and fluorescence intensities were analyzed on FNTA and nFCM, and used to plot a linear regression with MFI on Y-axis and number of AF488-labeled antibodies on X-axis in log_10_ scale (provided example is for anti-CD9-AF488 antibody used in COLO sEV experiments). Number of antibody equivalents per bead was assigned separately with plate reader by measuring serially diluted FS beads against each antibody used in this study. Data is shown as the mean ±SD of at least three independent experiments.

For comparison of instruments’ performances in fluorescence mode, fluorescent polystyrene FS beads of three different sizes (∼100, 40, and 20 nm) and brightness levels were used. Firstly, the beads were assigned the ERF value, by measuring them in the microplate reader against the serial dilutions of AF488-labeled antibodies used in the subsequent experiments (Supplementary file 1). Thus, the FS fluorescence intensity was expressed as the number of equivalent antibody molecules per bead. Due to the differences in F:P ratios, the assignment had to be done separately for each antibody type and production lot used in this study. Correction factors were devised to compensate for slight differences in the excitation and collection efficiencies of optics in microplate reader (excitation 485 nm - emission 535/25 nm bandpass), FNTA (excitation 488 nm – emission 500 nm longpass), and nFCM (excitation 488 nm – emission 525/40 nm bandpass). Marginal mismatching in excitation/emission spectra of the FS and AF488 was also taken into consideration. Thus, after ERF assignment was done by microplate reader, correction factor of ×1.17 was applied to the number of assigned antibodies for FNTA, and ×1.04 for nFCM (Figure S1, Supplementary files 1 and 2).

MFI of the beads measured on either FNTA or nFCM was converted to log_10_ scale and linear regression was plotted against the assigned ERF values, expressed in log_10_ as well (Figure 1C). Since the fluorescence intensity is not a native parameter of FNTA, it was calculated by multiplying *Area* and *Mean Intensity* of each detected event in the FCS file, followed by calculating the MFI of the whole population (Supplementary file 3). As shown in the representative linear regression plotted for anti-CD9 antibody (Figure 1C), nFCM displayed higher linearity between the data points, and better resolution, evidenced by a steeper slope. Since antibody assignment had to be done for each antibody type and production lot, individual linear regressions and equations were plotted and used for ERF conversions in the subsequent experiments.

### Optimizing NTA settings to define analytical acceptance criteria

When specific post-acquisition parameters (e.g., minimum and maximum particle area and brightness) are used in NTA measurements (either in scatter or fluorescence mode), the particles fitting the given criteria are detected and reported in the total particle concentration (particles/mL). However, only a fraction of these particles is successfully traced to acquire additional information, such as particle size and brightness (scattered or fluorescent light), which is then extrapolated as representative for total particles. If the acquired T:P ratio is low, or inconsistent between the measurements, this can introduce biases in the analyses. The shift in the T:P ratio can be especially noticeable with fluorescent particles which are dimmer and more difficult to detect (and trace) than scattering events.

Thus, to define the T:P acceptance criteria, in order to maintain reliability between our acquisitions and downstream result interpretation, we analyzed commercially available FLuoEVs (fluorescent EVs with CD63-EGFP fusion protein) in both scatter and fluorescence mode, using variable Minimum Brightness thresholds, while keeping all the other parameters unchanged. The amount of EGFP molecules per single EV was assessed based on the linear regression equation of the MFI and antibody assignments of the FS beads. The assigned antibodies per bead were converted to AF488 equivalents (based on the F:P information of multiple different antibodies; Supplementary file 1). Then, to account for the spectral and intensity differences, the AF488 was converted to EGFP by applying the correction factor of ×1.66, as described in the Materials and Methods. As shown in Figure 2A, the number of particles and traces detected in the scatter mode scaled proportionally with the reduction of the brightness threshold level. Nevertheless, the T:P ratio maintained ∼35% at all times. The traces that were lost in the higher threshold levels were, as expected, the smaller EVs, evidenced by the increased median particle size.

**Figure 2.**
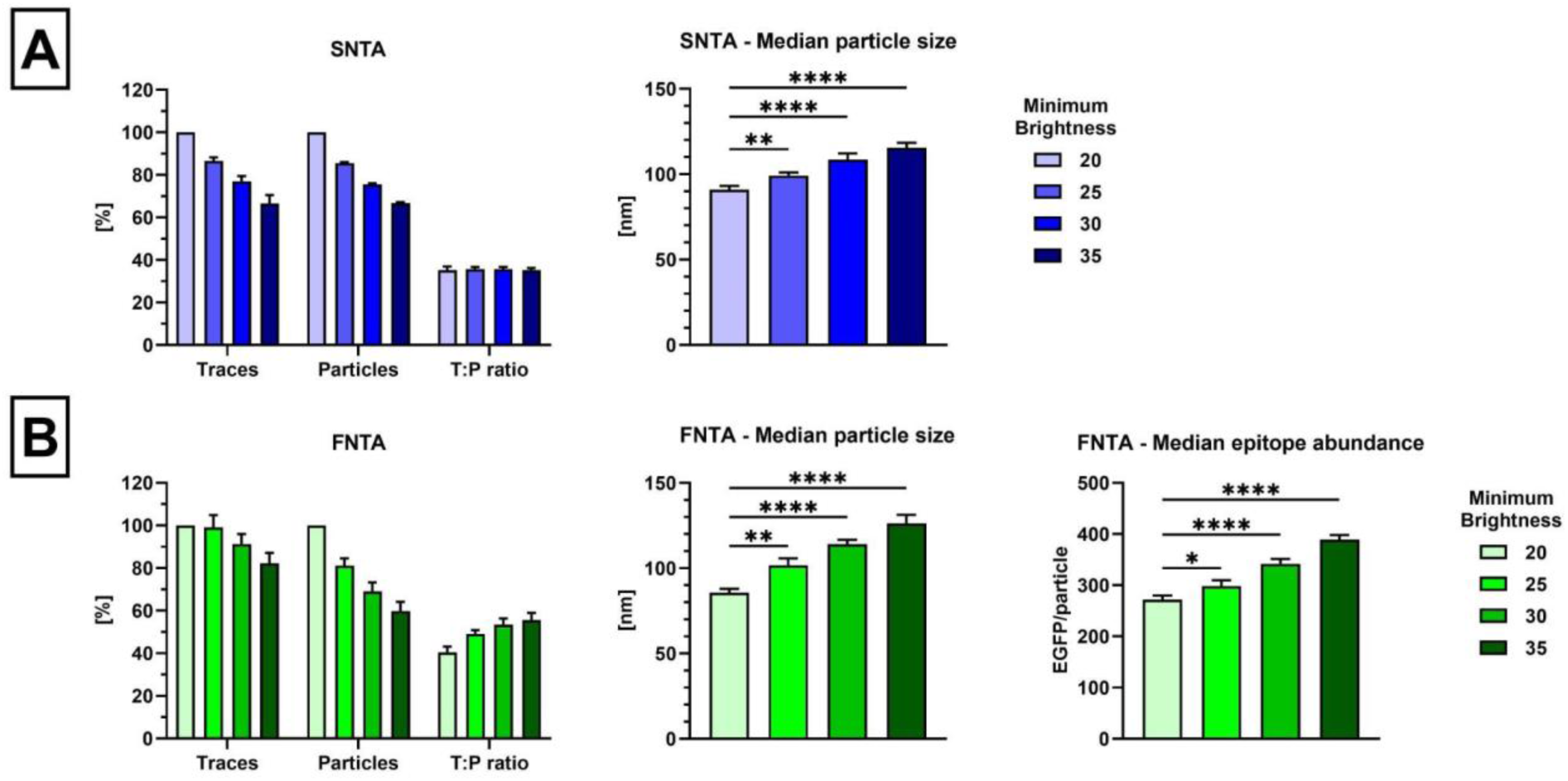
Effect of the minimum brightness levels on NTA analyses. Commercially available FLuoEVs (CD63-EGFP) were analyzed on NTA in both scatter and fluorescence mode (SNTA and FNTA). **(A)** By increasing the post-acquisition parameter, which defined minimum brightness of statistically relevant events, the number of detected particles and traces decreased, as expected. Nevertheless, the T:P ratio remained unchanged in SNTA, which means that the analyses kept the same level of representativeness for the detected particles. Even so, the traces became significantly larger in size at the higher threshold levels, indicating biased analyses. **(B)** In FNTA analyses, increasing the minimum brightness had a similar negative effect on the number of total detected particles. However, the number of traces did not change drastically. Thus, the T:P ratio improved, providing more representative analysis among the total detected particles. The tradeoff was that the median particle size and epitope abundance significantly increased with each threshold level. Ordinary one-way ANOVA with Dunnett’s multiple comparisons test was used for statistical analyses.

When FLuoEVs were analyzed in the fluorescence mode (FNTA, Figure 2B) with different Minimum Brightness thresholds, the number of traces was overall more stable, with only ∼18% loss at the highest threshold level. The number of total detected particles, on the other hand, showed a steeper decline. This consequently produced a higher T:P ratio (40% >> 56%), providing more representative results (i.e., estimation of particle size and epitope abundance) with respect to the whole population. However, this also created a bias in the traces, introducing the preference for significantly bigger and brighter particles (Figure 5B). Therefore, in order to acquire more representative data of the whole particle population, the tradeoff needs to be made by excluding smaller and dimmer particles from the analysis. With these experiments, we defined an acceptable T:P ratio to be at least 35%, for both scatter and fluorescence analyses, as it provided the optimal balance between the gain in data reliability and the loss of sensitivity. This strategy was applied and verified in the subsequent experiments with antibody-stained EVs, where the particles were much dimmer and more difficult to trace, compared to the scattering events or commercial FLuoEVs, thus yielding a very low T:P ratio (<15%). Such conditions urged us to use a variable Minimum Brightness threshold throughout the different measurements, in order to satisfy the acceptance criteria (example with antibody-stained EVs provided in the Figure S3). This is likely just the limitation of the current NTA software version. Future updates will probably bring significant improvements to the T:P ratio, even for the dimmest of the events, boosting the instrument’s sensitivity, resolution, and analytical reliability.

### Analysis of size, concentration, and epitope abundance on small and medium-sized EVs

After the initial comparison with synthetic nanoparticles and setting of the standardized protocols for fluorescence measurement expressed in ERF, purified small EVs from COLO cell line were labeled with either anti-CD9 or anti-CD63 antibodies and analyzed in both scatter and fluorescence mode on NTA and nFCM. In a scatter mode, particle concentration was comparable between the two instruments, while particle size appeared larger on SNTA (Figure 3A). Similar discordance in size estimation between SNTA and nFCM was reported before [26,27]. The results may vary due to the differences in the way each instrument calculates the particle size. NTA relies on the Brownian motion, while nFCM infers the size from the side scattered light. The latter is calibrated using silica beads, which still have a higher refractive index and scatter more light than EVs of the same size [16,28,29]. This might explain the comparable results between SNTA and nFCM for SiNPs in Figure 1, and a slight discrepancy in EV size estimation later on.

**Figure 3.**
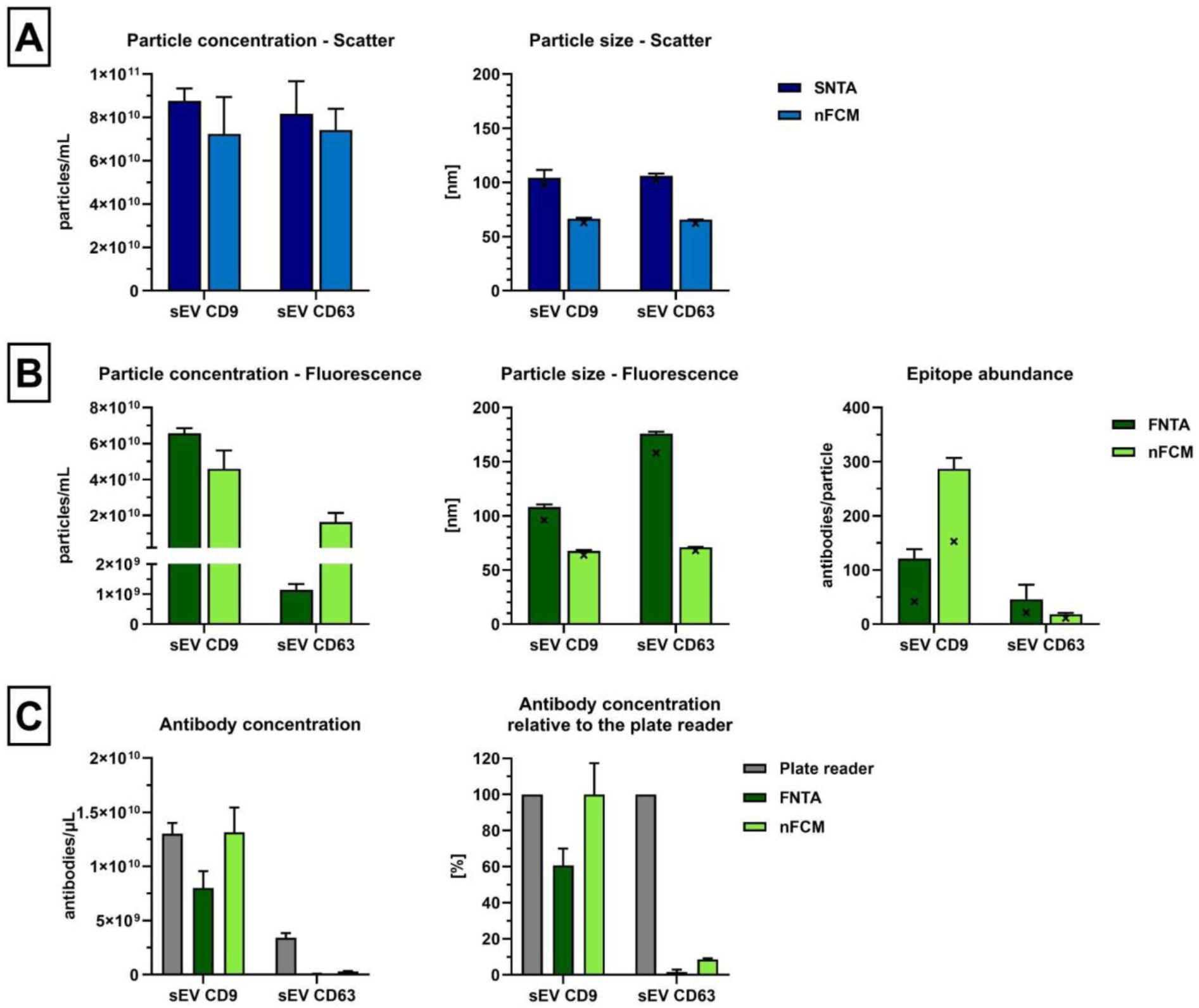
COLO sEVs labeled with anti-CD9 and anti-CD63 antibodies. COLO sEVs labeled with anti-CD9-AF488 and anti-CD63-AF488 were analyzed in both **(A)** scatter and **(B)** fluorescence mode on NTA and nFCM. Particle size is showing the average and median (bar and “x” symbol). Average and median (bar and “x” symbol) epitope abundance per particle were estimated using a linear regression formula plotted with the FS beads assigned with each antibody separately (Supplementary files 1, 2, and 3). **(C)** Antibody concentration per volume of sample was calculated from the average epitope abundance and the fluorescent particle concentration, and compared to the bulk fluorescence measurement performed in the plate reader. Data is shown as the mean ±SD of at least three independent experiments.

With the fluorescence measurements, the concentration of the CD9+ events was similar between the instruments, while CD63+ events were almost 15-fold higher on nFCM (Figure 3B and Supplementary file 4). The particle size for CD9+ events was comparable to the data obtained in a scatter mode, however, the CD63+ events appeared larger, especially on FNTA. Epitope abundance per single EV was estimated based on the linear regression equation obtained with FS beads and their respective ERF assignments. The average (and median) number of anti-CD9 antibodies bound per EV was significantly higher on nFCM than FNTA – 287 vs. 121 (median 153 vs. 42) antibodies/particle, respectively. The reason for such an imbalance in the epitope abundance for CD9+ events, in spite of comparable particle concentrations, was assumed to be sampling and analysis bias towards brighter particles. To confirm this hypothesis, we created the gating on nFCM which excluded just the top 1% or 5% of the brightest events. This in turn reduced the estimated antibody concentration by 18% and 37%, respectively, making the latter analysis more aligned with FNTA estimates (Figure S4). These observations highlighted the significance of the gating strategy and the impact it had on the final results. At the same time, when analyzing CD63+ events, nFCM was able to detect more particles with much lower average (and median) epitope abundance than FNTA – 18 vs. 46 (median 11 vs. 21) antibodies/particle, respectively. The increased brightness threshold for FNTA CD63 analyses, necessary to meet the T:P ratio criteria (see Materials and Methods), partly contributed to this outcome by limiting the analyses to mostly bigger particles displaying more epitopes (Figure 3B). The same samples were analyzed also in a plate reader, and the overall antibody concentration was calculated as described in Materials and Methods (Figure 3C and Supplementary file 4). This bulk measurement was compared with FNTA and nFCM readouts, and it showed that the latter enabled more accurate estimation of EV-bound anti-CD9 antibody concentration. As previously mentioned, and briefly demonstrated in the Figure S4, the reason for this may lie in the CD9 epitope stoichiometry being greatly unbalanced across the whole EV population, with the largest portion of bound antibodies presented on very few brightest events. On the other hand, concentration of anti-CD63 antibodies in the sample was greatly underestimated by both nFCM and FNTA (Figure 3C and Supplementary file 4). This meant that most of the CD63+ events are still under the limit of detection (LoD), due to the low CD63 antigen expression per single small EV. Similar observations were made in a previous study in which Western blot bulk analyses showed higher CD63 marker expression in plasma EVs purified by SEC, even though single particle measurements indicated otherwise [30]. Contrary to CD9, CD63+ events appeared to have more homogeneous epitope stoichiometry, since they did not have any extremely bright events in the whole population, evidenced by a very low average antibodies/particle, closer to the median. As previously reported, the overall detectability of the signal is also affected by the fluorophore brightness and F:P ratio of the antibody used [31]. Considering that AF488 is much dimmer than some alternative fluorophores, e.g., PE (∼25x brighter than AF488), even the high F:P ratio might not be enough for successful detection of particles like CD63+ COLO sEVs.

Altogether, our results highlight the importance of gating strategy, antigen expression level, fluorophore brightness, and F:P ratio for the inclusion or exclusion of certain fluorescent events from the analysis, thus affecting the total antibody detection by FNTA or nFCM. In such a scenario, bulk measurement of total antibody signal by a plate reader can serve as a good normalizer in order to appreciate to which extent the single EV characterization represents the detection of total EV-bound antibodies.

We hypothesized that by increasing the particle size, the CD63 epitope abundance would increase as well, raising the events’ brightness above the LoD, and enabling more reliable and inclusive single particle analyses than with sEVs. To test this hypothesis, we analyzed medium size EVs (mEVs >150 nm) from the same COLO cell line labeled with the same anti-CD63 antibody. When comparing the fluorescence analyses between the FNTA and nFCM, the results were relatively in line with the initial observations made with CD63+ sEVs. With nFCM, we detected more particles with lower epitope abundance, while FNTA measured mostly larger particles with higher epitope abundance (Figure 4A). This time, the discrepancy in particle concentration between the FNTA and nFCM was not as prominent as before (∼2-fold with CD63+ mEVs vs. ∼15-fold with CD63+ sEVs), indicating that the events became bright enough for improved FNTA detection. The apparent particle size was unexpectedly small on nFCM (average of ∼100 nm). To a certain degree, we can suspect that some of the additional particles detected by nFCM were also smaller, causing the shift in the size estimate. On the other hand, we acknowledged that using silica beads as a reference material for EV size calibration may be suboptimal. The average epitope abundance was now increased, compared to the CD63+ sEVs (93 antibodies/particle for FNTA and 24 antibodies/particle for nFCM), and overall estimated antibody concentration in the sample drastically improved (Figure 4B). Similar to CD9+ sEVs before, we observed that the unbalanced CD63 epitope stoichiometry in mEVs now created an analytical bias - the fewer brightest events, carrying the majority of antibodies, skewed the antibody concentration results, this time in favor of FNTA, even though nFCM demonstrated higher sensitivity by detecting more particles. Nevertheless, it was still evident that plenty of antibodies were left undetected on both instruments. At least in part this can be attributed to the unbound and unwashed antibodies that were not detected in single particle measurements, but are still present in bulk analyses, despite the intensive washing steps. Even with this persisting limitation, we were able to prove our hypothesis that larger particles facilitate detection of underexpressed markers. Detailed results can be found in the supplementary file 4.

**Figure 4.**
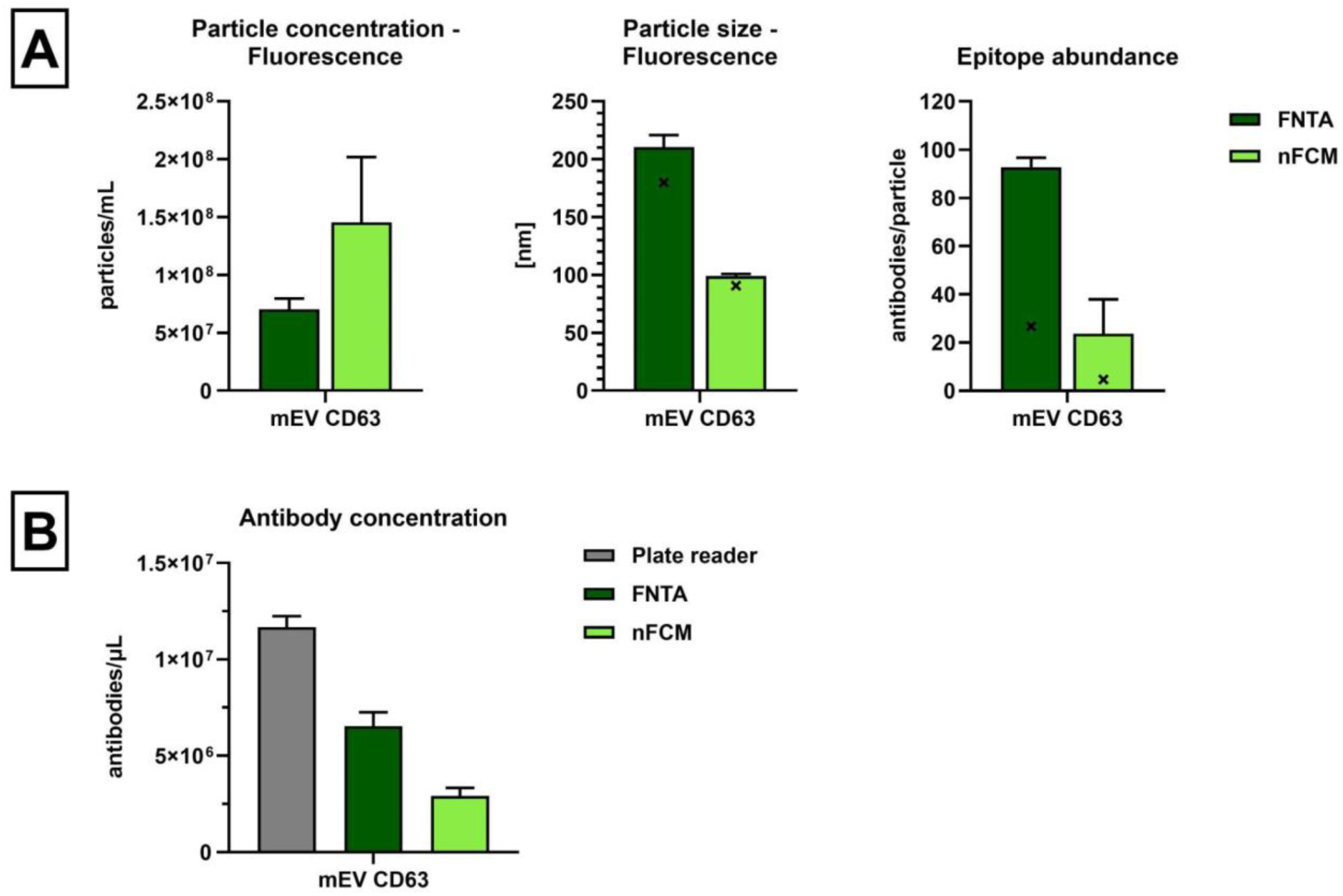
COLO mEVs labeled with anti-CD63 antibodies. **(A)** COLO mEVs labeled with anti-CD63-AF488 were analyzed on FNTA and nFCM. Particle size is showing the average and median (bar and “x” symbol). Average and median (bar and “x” symbol) epitope abundance per particle were estimated using a linear regression formula plotted with the FS beads assigned with the anti-CD63 antibody (Supplementary files 1, 2, and 3). **(B)** Antibody concentration per volume of sample was calculated from the average epitope abundance and the fluorescent particle concentration, and compared to the bulk fluorescence measurement performed in the plate reader. Data is shown as the mean ±SD of at least three independent experiments.

### Estimation of epitope abundance on the small EVs from complex biofluid samples

We proceeded with our experiments by analyzing more complex samples – EVs purified from plasma using SEC. The particles in such a sample display great heterogeneity due to the presence of EVs from multiple cell and tissue sources, as well as the abundance of larger lipoproteins (LPs) that co-elute with EVs. We targeted CD9 and CD63 to assess the broad range of EVs among the total EV and non-EV particles, while CD41 and glycophorin A (GYPA) were meant for detection of two most abundant subpopulations derived from platelets and red blood cells, respectively. As expected from such a polydisperse sample, we once again saw the higher particle concentration on SNTA compared to nFCM (Figure 5A). The fluorescence analyses proved to be much more challenging with this sample as the LP:EV ratio was in favor of the former, yielding an extremely low number of events labeled with antibodies against general or blood-lineage specific markers [32–34]. Still, nFCM was able to detect significantly more particles, which were smaller and expressed fewer epitopes per single particle. Our measurements are in line with the distribution of EVs in plasma that is reported to be tilted towards small, nanosized vesicles [35]. The lowest epitope abundance measured on nFCM was for GYPA+ EVs, with an average of 11 (median of 5) antibodies/particle. The highest average epitope abundance measured on nFCM was with CD41, at 130 antibodies/particle, while the highest median epitope abundance was detected with CD63, showing 17 antibodies/particle (Figure 5B and Supplementary file 4). In fact, we found that the CD9 and CD63 average epitope abundance estimations made with nFCM, were in fine agreement with earlier studies performed with qSMLM [14]. The concordance extended also to the measured average particle size, even if slightly smaller on nFCM. The reason for measuring apparently smaller size on nFCM than qSMLM is likely the same as mentioned before, i.e., due to the usage of silica beads as a reference material for the scattering profile of EVs.

**Figure 5.**
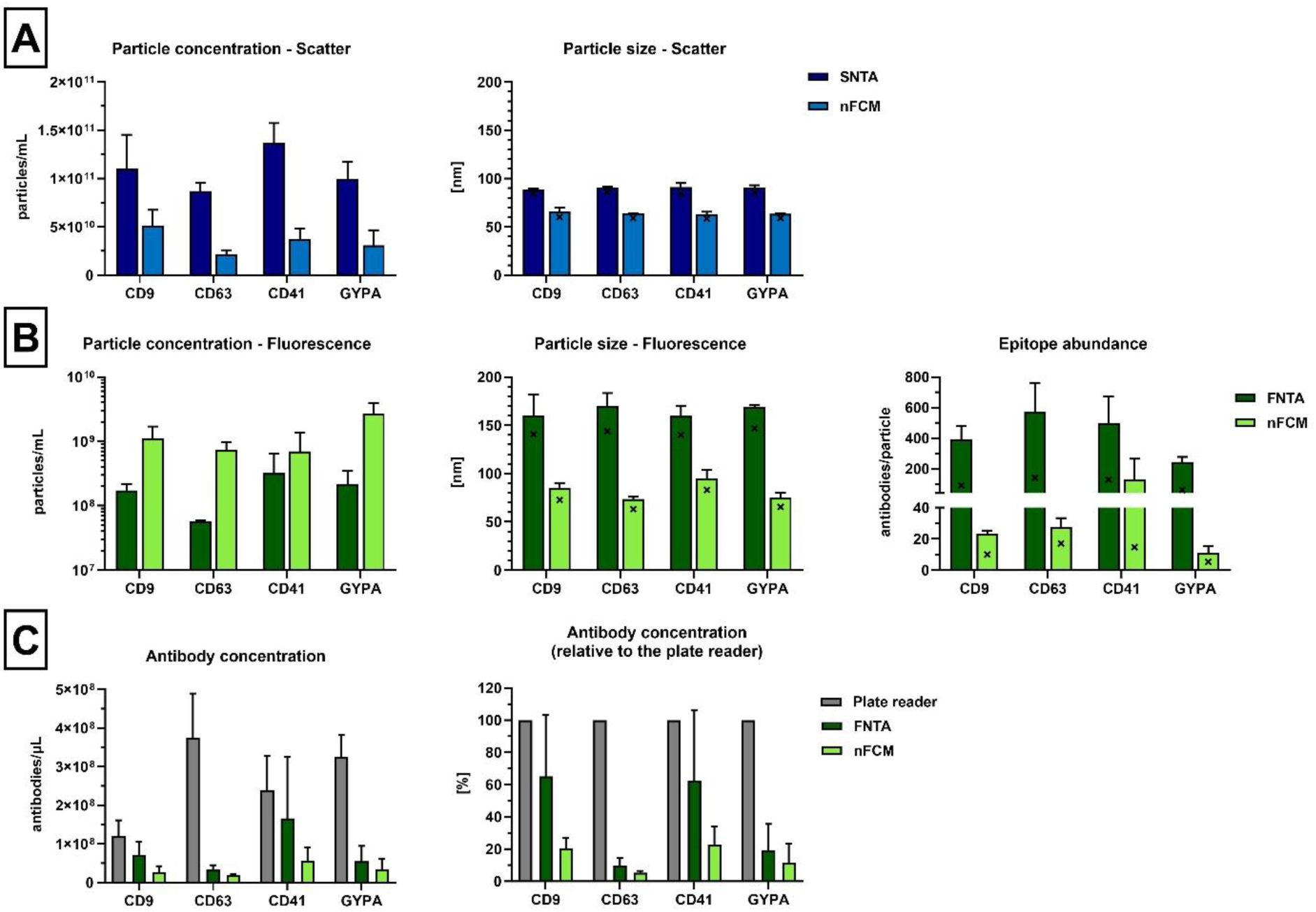
Plasma EVs labeled with antibodies against general and blood-lineage specific markers. SEC-purified plasma EVs labeled with anti-CD9, anti-CD63, anti-CD41, and anti-GYPA antibodies (AF488 conjugates) were analyzed in both **(A)** scatter and **(B)** fluorescence mode on NTA and nFCM. Note the log_10_ scale for the fluorescent particle concentration. Particle size is showing the average and median (bar and “x” symbol). Average and median (bar and “x” symbol) epitope abundance per particle were estimated using linear regression formula plotted with the FS beads assigned with each antibody separately (Supplementary files 1, 2, and 3). **(C)** Antibody concentration per volume of sample was calculated from the average epitope abundance and the fluorescent particle concentration, and compared to the bulk fluorescence measurement performed in the plate reader. Data is shown as the mean ±SD of at least three independent experiments.

Due to the sensitivity of the FNTA, the majority of detected particles were larger EVs with higher epitope abundance. FNTA struggled to detect and maintain the trace length for many of the events throughout the acquisition, requiring a higher Minimal Brightness threshold in order to maintain T:P ratio ≥35%, comparable to the scatter measurements (see Materials and Methods). As the result, particles included in the statistical analyses were much larger and brighter, but still maintained similar marker expression trend as observed with nFCM – the lowest was GYPA, with an average of 245 (median of 61) antibodies/particle, and the highest was CD63, with an average of 574 (median of 140) antibodies/particle. For most of the analyzed samples, both FNTA and nFCM were detecting 5-20% of the total antibodies compared to the bulk plate reader estimates (Figure 5C and Supplementary file 4). However, with the FNTA’s biased analyses of mostly brighter events, the estimated antibody concentration was significantly higher in case of CD9 and CD41 markers, reaching more than 60% of the bulk estimates. Once more, we observed that the biased analysis and a non-Gaussian distribution of certain epitopes are the main reason for such highly concordant results between FNTA single particle analysis and microplate reader bulk measurements. With the exclusion of just 1% or 5% of the top brightest events, the number of detected antibodies in FNTA dropped by 22% or 36%, respectively (Figure S5).

### Approximating the instruments’ limits of quantification

Since nFCM produced better linearity and resolution than FNTA for fluorescence analyses, we provided the overview of their respective accuracies (portrayed in residual plots), and approximated their LoQ. Unlike the LoD, which is defined as the lowest detectable MFI value above the background, here, the LoQ refers to the MFI signal which instruments can measure with greater reproducibility. Using the F:P ratio of multiple antibodies (information provided by the manufacturer), and appropriate correction factors (as described in Material and Methods), we converted ERF of FS beads from antibodies/bead to AF488/bead (Table 3). The linear regression was then plotted using measured MFI values and assigned AF488 values of FS beads (log-transformed), and the equations were used to back-calculate the AF488/bead. Following formula was used to calculate the residuals for both FNTA and nFCM: 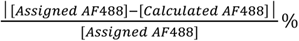. Due to the apparently poor detection of the dimmest FS beads, the residuals were quite high in the lower ranges, more so for FNTA than for nFCM. Based on this, we used FS20 beads as our reference for the lowest quantifiable MFI on both instruments. We then calculated an “assumed AF488/bead” value that would ensure better linearity (with residuals <15%) for the same measured MFI. This value served as an approximate LoQ, estimated at ∼115 AF488 molecules for FNTA and ∼75 AF488 molecules for nFCM (Table 3). This further translates to ∼25 and ∼16 antibodies per particle, respectively, although, subjected to a high variation depending on the individual antibody’s F:P ratio. For example, anti-GYPA-AF488 antibody, with the F:P = 5, provided better detection of dimmer events, than anti-CD41-AF488, with F:P = 2.9 (Figure 4B). It is worth noting that the LoD reached levels as low as couple of antibodies in most of the measurements.

**Table 3.**
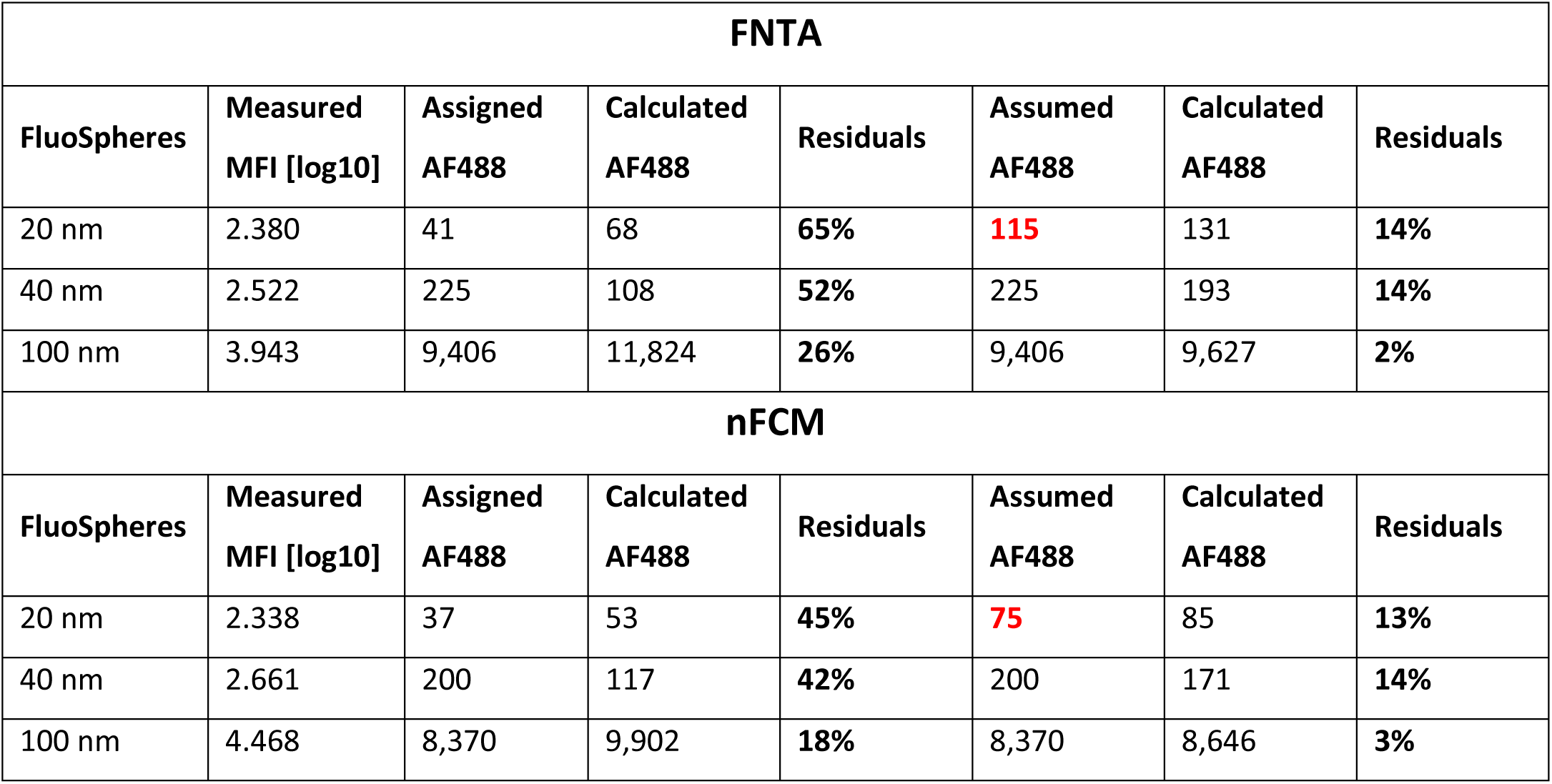
Residuals in linear regression analysis with FNTA and nFCM, and approximate instruments’ LoQ.

| **FNTA** |  |  |  |  |  |  |  |
| --- | --- | --- | --- | --- | --- | --- | --- |
| **FluoSpheres** | **Measured**<br>**MFI [log10]** | **Assigned**<br>**AF488** | **Calculated**<br>**AF488** | **Residuals** | **Assumed**<br>**AF488** | **Calculated**<br>**AF488** | **Residuals** |
| 20 nm | 2.380 | 41 | 68 | **65%** | **115** | 131 | **14%** |
| 40 nm | 2.522 | 225 | 108 | **52%** | 225 | 193 | **14%** |
| 100 nm | 3.943 | 9,406 | 11,824 | **26%** | 9,406 | 9,627 | **2%** |
| **nFCM** |  |  |  |  |  |  |  |
| **FluoSpheres** | **Measured**<br>**MFI [log10]** | **Assigned**<br>**AF488** | **Calculated**<br>**AF488** | **Residuals** | **Assumed**<br>**AF488** | **Calculated**<br>**AF488** | **Residuals** |
| 20 nm | 2.338 | 37 | 53 | **45%** | **75** | 85 | **13%** |
| 40 nm | 2.661 | 200 | 117 | **42%** | 200 | 171 | **14%** |
| 100 nm | 4.468 | 8,370 | 9,902 | **18%** | 8,370 | 8,646 | **3%** |

## DISCUSSION

In this work we have compared two single particle analysis platforms which are commonly used for EV analysis – NTA and nFCM. We evaluated their performance in both scatter and fluorescence mode using synthetic nanoparticles (silica and fluorescent polystyrene beads of different sizes and fluorescence intensities), and EVs from simple (cell culture media) and complex biofluids (human blood plasma).

Overall, when the samples were monodisperse, containing relatively small particles (∼100 nm), both instruments assessed the concentration equally well. However, with the measurements of polydisperse samples, we observed that nFCM results were more in line with the expected values (according to the polydisperse silica beads mix), since larger events might have caused the overestimation on NTA, as reported earlier [26]. We acknowledge that this was likely a limitation of using a single instrument setting on NTA for analyzing a wide range of particles in our experiments. Furthermore, in most of the EV measurements, NTA reported larger particle size compared to nFCM. To some degree, this can be caused by the intrinsic differences in the way these instruments perform size measurements – NTA calculates the particle size from the Brownian motion, while nFCM relies on the particle’s refractive index and side scattering light. The latter is calibrated using silica beads as a reference material, which introduces a level of uncertainty, because solid silica beads scatter more light than EVs [28,29]. The disparity between NTA and nFCM size measurements was even more pronounced in fluorescence mode, since FNTA biased larger EVs with concomitantly higher marker abundance.

In fluorescence mode, both instruments were struggling to detect the dimmest events, although nFCM was more sensitive, as demonstrated by greater linearity and resolution produced with FS beads of three different levels of fluorescence intensity. The residuals in lower ranges were still significant as we were operating on the very limit of detection for both instruments. Nevertheless, our method still enables a higher level of quantification, standardization, and cross-platform comparability than the relative fluorescence units (which are specific to each instrument), or just purely digital quantification (positive vs. negative particles in fluorescence measurement). In fact, the level of accuracy of our standardization approach seems to outperform current state-of-the-art calibration techniques with conventional flow cytometry MESF calibrators [18]. The recent work of Lozano-Andrés et al. showed that despite the high linearity and complementarity between different batches of beads for the similar quantitative range, these calibrators still failed to project a comparable level of accuracy and confidence in the fluorescence range of EVs. Reported variation in MESF estimates, based on the used flow cytometry calibrators, reached 157% for EVs stained with anti-CD9-PE, while analyzing true residuals in the EV range was virtually impossible since it was at least 2 orders of magnitude below the quantification range of the beads. Having a fluorophore that is ∼24x brighter than AF488 (brightness of PE = 1,607,200 M^-1^cm^-1^ vs. AF488 = 67,160 M^-1^cm^-1^), did not seem to help in mitigating the circumstances. All of the aforementioned observations demonstrate the limitations of using big and bright beads from flow cytometry to extrapolate and calibrate the low levels of fluorescence intensities of EVs. Together with the fact that the flow cytometry beads are incompatible with instruments such as FNTA and nFCM, this makes a compelling argument for the adoption of our fluorescence standardization method with FS beads.

Hard-dyed beads are generally not recommended for fluorescence calibration, due to the high spectral mismatching (usually with multi-peak emissions) with respect to the fluorophores used to stain EVs [16,19]. However, the yellow-green FS beads show a single-peak emission spectrum, highly concordant to that of AF488, making it a suitable candidate for such application. Standardized production, which ensures low polydispersity index (6-14% for FS batches used in our experiments), and accurate bead concentration (derived from the dry weight of the beads, the density of polystyrene, and the bead diameter), further supports the suitability of FS beads for ERF assignment. Indeed, a similar strategy was reported before, using soluble AF488 and yellow-green FS beads for ERF assignment [36,37]. In the former of the two cited articles, the authors demonstrated applicability of hard-dyed beads in standardizing the fluorescent single particle analyses by measuring highly linear fluorescence intensity between three FS bead populations (∼40, 100, and 200 nm). Since we used one set of smaller and dimmer beads (20 nm) for our experiments, it explains the loss of linearity in the lower ranges, which we discussed in the previous paragraph, although it provided a better coverage of a true EV quantification range. Furthermore, our approach allowed for direct assessment of antibodies (not fluorophore molecules) per particle, consequently getting closer to an estimation of epitope abundance.

We were also able to approximate the instruments’ LoQ at about 115 and 75 AF488 equivalents for FNTA and nFCM, respectively. This can be translated to ∼5 and ∼3 PE molecules, respectively, with the consideration of above-mentioned brightness disparity (PE being 24x brighter than AF488). The prospective LoD is likely lower. Even with such respectable sensitivity levels, we observed that many fluorescent events went undetected (by comparing FNTA, nFCM, and plate reader data). We cannot entirely rule out that some of the signal detected by the plate reader originated from the unbound antibodies that might have remained in the sample even after rigorous washing, although our dye+PBS controls showed complete antibody removal (data not shown). Apart from that, the lack of detection of many fluorescent events by FNTA and nFCM was likely influenced by the low antigen expression level per single EV, as well as the fluorophore brightness and F:P ratio of the antibody used (discussed below). In addition to the sensitivity limitations that can create a bias towards brighter particles, gating strategy appeared also as a factor that can have a significant impact on the detectability of events, in this case on both sides of the extremes – dimmest and brightest particles. The final result, comparing single particle analyses and bulk measurements, was particularly influenced by the level of inclusiveness of the brightest events, which hauled majority of the bound antibodies (demonstrated also in the Figures S4 and S5). By focusing only on the fewer brightest events, we were able to detect most of the bound antibodies in the sample, and acquire results that had highest complementarity with the plate reader bulk analyses data. Such analytical paradox may instigate the polemics regarding the race for development, and finally utility, of the most sensitive instrument for EV analyses. The argument in favor would be that the higher sensitivity conveys more informative assessment of EVs, with insights into their true heterogeneity and marker stoichiometry. We can consider that, although heterogeneous in size, EVs distribution in human plasma is biased towards smaller vesicles (smaller than 100 nm) [35]. High sensitivity capabilities are particularly advantageous for diagnostic application of EVs, where, in spite of the increased tumor-EV production rate in circulation of patients (5-15x higher than in healthy individual), the disease biomarker abundance per single EV is significantly lower (e.g., 16 CA19-9 molecules/EV in cancer patients vs. 31 in healthy individuals) [15]. In this case, EV analytical instruments lacking sensitivity would not be able to detect the tumor-derived EVs.

The previous paragraph highlights three additional factors that, together with an instrument’s gating strategy and overall sensitivity, shape the final result of EV analyses on a single particle level. These factors are marker abundance per single EV, fluorophore brightness, and antibody’s F:P ratio. The fact that the number of conjugated fluorophores per antibody can have a significant influence on the EV detectability was recently reported in the work of Weiss et. al. [31]. The most prominent case where all these factors might produce negative synergistic effect, with the highest impact on the future of EV diagnostics, was indeed the detection of different subpopulations of circulating EVs. Based on our results, we made an illustrative model showing the interplay of instrument’s sensitivity and gating, marker abundance, and F:P ratio of the antibody used to detect the EV displayed protein marker of interest (Figure S6). In the case where the average marker expression was high enough per single EV (e.g., CD41), both FNTA and nFCM were able to detect a fair number of EVs, in spite of the low F:P ratio of the antibody (∼3 AF488 per anti-CD41 antibody) and FNTA’s sensitivity level that favors brighter events even with the lowest threshold setting (minimum brightness set at 20). On the contrary, analyzing a marker with low abundance per single EV (e.g., GYPA) proved to be much more challenging for FNTA, especially with a stricter gating strategy, ensuring the optimal traceability of events (minimum brightness set to 25-30, instead 20). At the same time, nFCM benefited from the usage of an antibody with higher F:P ratio (∼5 AF488 per anti-GYPA antibody) as it allowed detection of numerous events with extremely low epitope abundance. Such analyses also raise interesting points on plasma EV biology, and provide deeper understanding of the true EV heterogeneity, by bringing into perspective the dynamic expression and distribution of different EV markers. This information can feed the future *in silico* models, which could aid in EV analytics, boost the EV biomarker discovery and validation, and facilitate the development of EV-based liquid biopsies.

Considering the previous notions on the biology of plasma EVs, we further compared the scientific consensus in the literature with our observations of the total plasma EV concentration, and the subpopulation heterogeneity. By looking at the average positivity for all of the markers and normalizing it per volume of input biofluid, we assessed the EV concentration with FNTA to be ∼1.9×10^9^ EVs/mL of plasma, while with nFCM it was ∼1.4×10^10^ EVs/mL of plasma, which was more in line with the expected estimates of ∼2×10^10^ EVs/mL of healthy-donor plasma [33,34]. Of course, we also keep in consideration that SEC was used for EV isolation from plasma prior to analysis, and that the recovery of EVs from a complex fluid can have variable efficiency that is also dependent on the donor sample variability. Similar to the previous report [30], we observed with nFCM that the red blood cell (RBC)-derived EVs (GYPA+) were more abundant than those from platelets (CD41+). This may appear somewhat conflicting, since some other studies, as well as our FNTA results, indicate otherwise [34,38–41]. In certain instances, such a discrepancy could be explained by the extent of platelet activation in different experiments, which could create EV artifacts if the pre-cleared plasma still has some remaining platelets, or if serum is used instead. On the other side, from a physiological point of view, the EV-mediated material removal is suggested to be implicated in a reticulocyte-to-erythrocyte maturation [42], the continuous and high-turnover process, making the predominance of GYPA+ vesicles in human blood plausible. Another point of view would be that the stoichiometry and epitope abundance were not considered in the previous studies, leading to conclusion that platelet EVs make up the majority, while in fact, most of the RBC EVs are just undetectable, due to the low epitope abundance per single particle and the lacking analytical sensitivity, as discussed before. This presumption is well in agreement with the provided example, in which an Apogee A60 instrument, calibrated for the measurement of EVs in the size range of 160-1,000 nm, was used for the analyses of platelet- and RBC-derived EVs in plasma [40]. Even though the measured relative proportion of these subpopulations is comparable to what we observed with FNTA (more platelet EVs than RBC EVs), their actual concentration appears to be ∼50x and ∼100x lower than FNTA estimates (or ∼130x and ∼1,300x lower than nFCM estimates), respectively. This suggests that the vast majority of plasma EVs were probably not detected within such a high calibration range (160-1,000 nm), and most likely, the RBC EVs took a bigger sacrifice in the analyses, due to the much lower GYPA abundance per single EV. In that case, it does not surprise that with progression in the sensitivity from lower to higher (Apogee A60 < FNTA ZetaView < nFCM NanoAnalyzer), not only that we see the change in the concentration of detected EVs, but also the change in the subpopulation proportions, i.e., RBC EVs becoming more abundant than platelet EVs.

Looking into the expression of the CD41 and GYPA markers on a cellular level, we can divulge and debate the non-randomness of molecular sorting during the EV biogenesis, which creates the marker stoichiometry suboptimal for EV detection. Apparently, one platelet of ∼2.5 µm has ∼40,000 CD41 molecules on its surface [43], while an average-sized RBC can display up to 1,000,000 GYPA copies [44,45]. This translates to an epitope density (marker occupancy per surface area of the cell membrane) of approximately 1 CD41 molecule per 490 nm^2^ and 1 GYPA molecule per 134 nm^2^ [46]. One could then justifiably expect that the GYPA expression would be almost 4x higher than CD41, per single EV of the same size. Yet, measurements show quite the opposite. The results of nFCM analyses show that CD41+ EVs have similar epitope density as their parental cells (∼1 molecule per 445 nm^2^; based on the average epitope abundance and average fluorescent particle size), while GYPA+ EVs with 1 molecule per 1,749 nm^2^ demonstrate significantly lower epitope density compared to their cell of origin. This may suggest the existence of an apparently non-stochastic process, with high level of order in cellular EV packing machinery, that leads to preferential depletion of GYPA from RBC EVs. On the opposite side, CD41 in platelet EVs appears to follow no particular sorting instructions. It is also plausible that sEVs detected in our study originate both from endosomal compartments and from plasma membrane budding of their parent cells. Two biogenesis pathways can result in distinct molecular composition of vesicles and may also be unequally favored in different cell types. For instance, platelets are reported to produce microvesicles (membrane originated) and exosomes (of endosomal origin), both displaying platelet antigens, but only latter being enriched in CD63 [39,47]. In reticulocytes, instead, the selective removal of proteins and enzymes during the erythrocyte formation is predominantly occurring through exosomes [48], possibly explaining their different membrane composition with respect to that of the parent cell. Of course, we have to make our interpretations cautiously due to some procedural and instrumental factors that might influence the epitope (and epitope density) readout, such as antibody’s affinity and avidity, or instrument’s inability to detect all of the EVs, hence, providing biased results. In spite of these limitations that could restrict direct comparison of EV and cellular epitope densities, certain relative comparisons could still be made among EVs alone. Deeper analyses of individual EVs (using FCS files) might reveal stoichiometric subpopulations within subpopulations, i.e., groups of EVs expressing the same marker but with different epitope densities. For example, looking at the CD9+ COLO sEVs on FNTA in the size range 40-100 nm and 100-200 nm, we could see that the smaller subpopulation has on average 100 antibodies/particle with epitope density of ∼570 nm^2^, while the larger subpopulation had an average of 135 antibodies/particle and density of ∼1,500 nm^2^.

Finally, having an absolute quantification of EV markers and understanding their stoichiometry is particularly important when assessing disease-associated EVs. In the study performed by Lennon et al. [15], mentioned in the previous paragraph, plasma EVs from healthy donors and pancreatic cancer patients were analyzed by qSMLM, revealing that, even though subpopulations of EGFR and CA19-9 increased in concentration in cancer patients, these EVs were larger and carried less markers, i.e., had drastically lower epitope density - 1 molecule per 131-242 nm^2^ in healthy samples vs. 707-730 nm^2^ in disease samples. Such a shift in stoichiometry within the subpopulation could render them undetected by the instruments with low sensitivity, or at least create a bias in the analyses towards bigger EVs or different stoichiometric subpopulations.

## CONCLUSIONS

Our study introduces a novel method for standardizing fluorescence quantification and facilitating comparison across different single-particle analysis instruments, such as FNTA and nFCM. This approach enables deeper characterization of EV subpopulations, and better understanding of analytical challenges.

We further highlight the significance of instrument’s gating strategy and sensitivity, fluorophore brightness, and antibody’s F:P ratio in shaping the final result of single EV analyses, and propose in-parallel bulk fluorescence measurements as a reliable tool for data normalization.

We also evidence the opportunity and feasibility of gating strategies and software adjustments to boost and balance the instrument sensitivity and analytical reliability, both paving the way towards more robust EV biomarker detection.

Overall, our work provides a valuable contribution to the ongoing development and optimization of EV analytical platforms and future EV diagnostic tools.

## ACKNOWLEDGMENTS

This work was supported by following funding programs: Tallinn University Research Fund (project number TF7320), European Regional Development Fund Enterprise Estonia’s Applied Research Program under the grant agreement number 2014-2020.4.02.21-0398 (EVREM).

