## Supplementary figures for "Quantitative fluorescent nanoparticle tracking analysis and nano-flow cytometry enable advanced characterization of single extracellular vesicles"

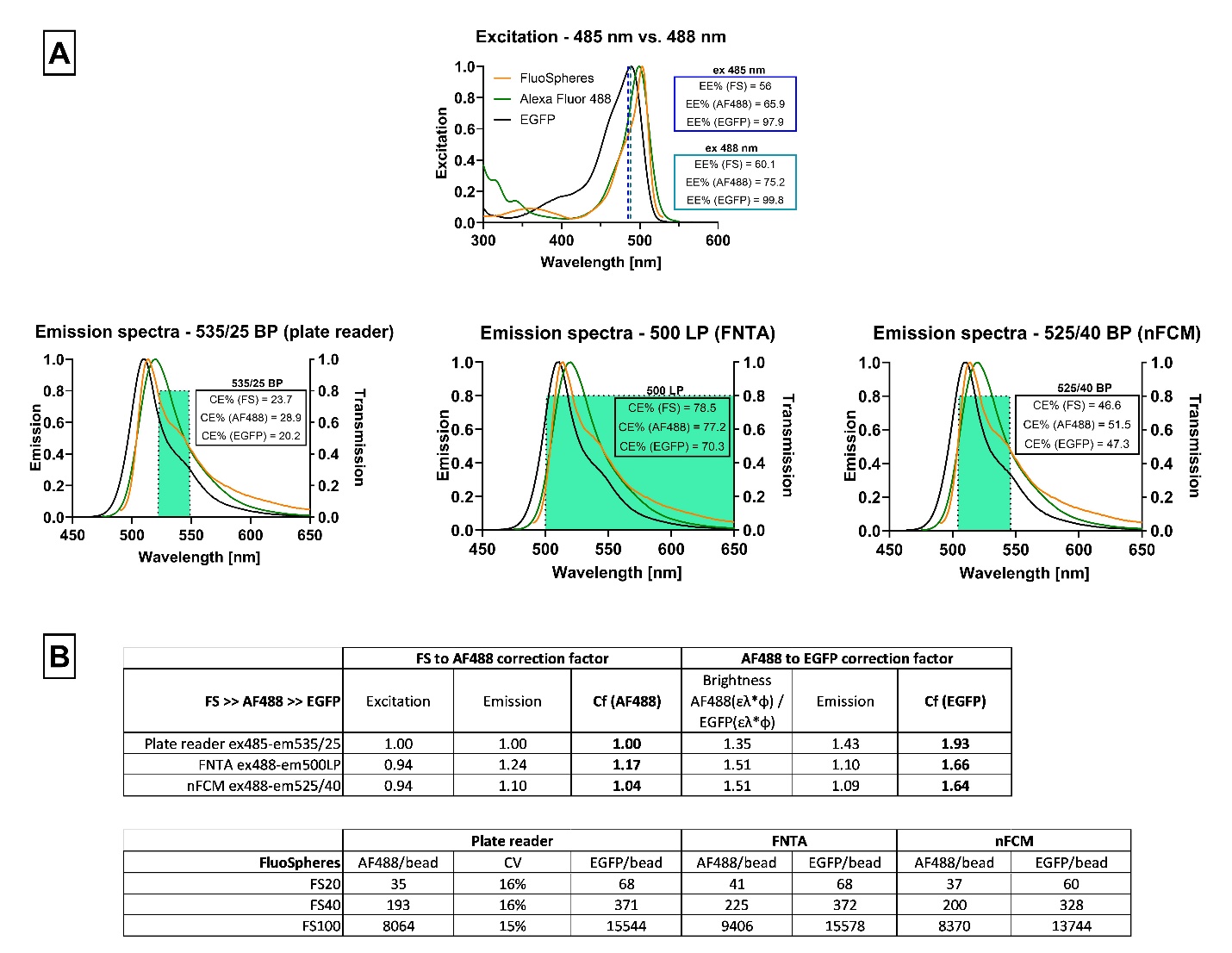

**Figure S1. Excitation/collection efficiencies and correction factors for different fluorophores.** **(A)** Excitation efficiencies (EE%) with 485 nm and 488 nm laser lines, and emission spectra are shown for FS beads, AF488, and EGFP. Highlighted (green) area represents the collection efficiency (CE%) for optical filters in plate reader (535/25 nm bandpass), FNTA (500 nm long-pass), and nFCM (525/40 nm bandpass). Transmission efficiency was assumed to be uniform in all three instruments (T = 80%). **(B)** The correction factors were devised from the efficiencies of the given lasers and filters, which was then applied for converting FS MFI signal to the number of antibodies or AF488 molecules, and further for converting AF488 to EGFP molecules.

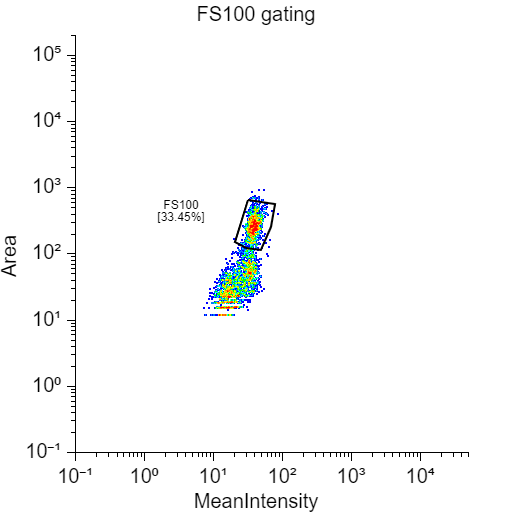

**Figure S2. Gating strategy for FS100 beads.** Since the FS100 beads were too bright for the FNTA SOP that was used with EVs, the gating was required in order to isolate only those beads that were in the focal plane.

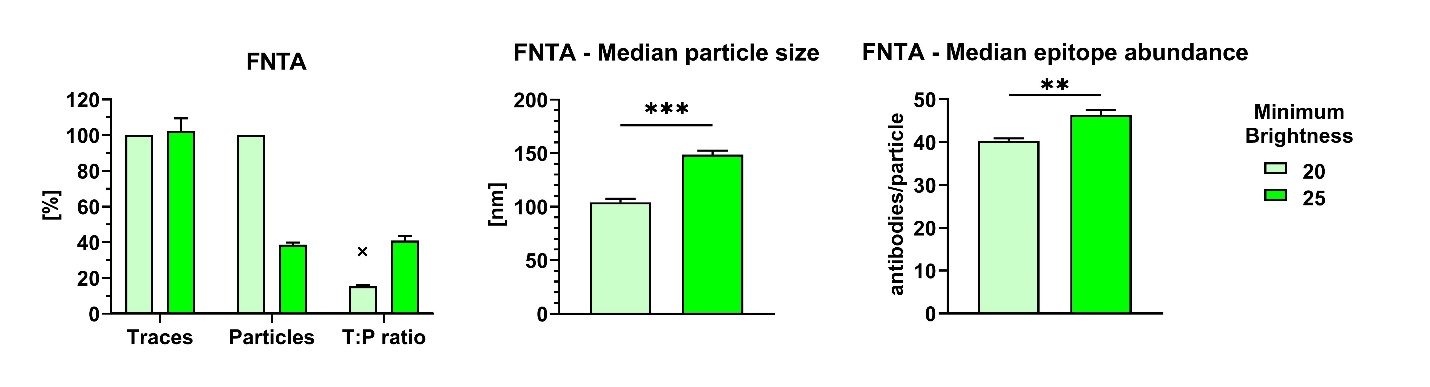

**Figure S3. Effect of the minimum brightness levels on FNTA analyses of antibody-stained EVs**. For analyses of antibody-stained EVs (provided example is for plasma EVs stained with anti-GYPA-AF488 antibody)**,** switching to a higher threshold level (from 20 to 25) did not affect the number of traces, however, it reduced significantly the number of detected particles, since the sample contained many dim events, which NTA software was not able to trace. This in turn produced a more noticeable shift in T:P (15%>>41%), compared to the FLuoEV measurements (Figure 2), providing more reliable and representative statistics (relative to the whole particle population). The tradeoff was a stronger bias towards bigger and brighter particles. Symbol “x” shows the expected T:P ratio according to the analyses in the scatter mode. Unpaired two-tailed t test was used for statistical analyses.

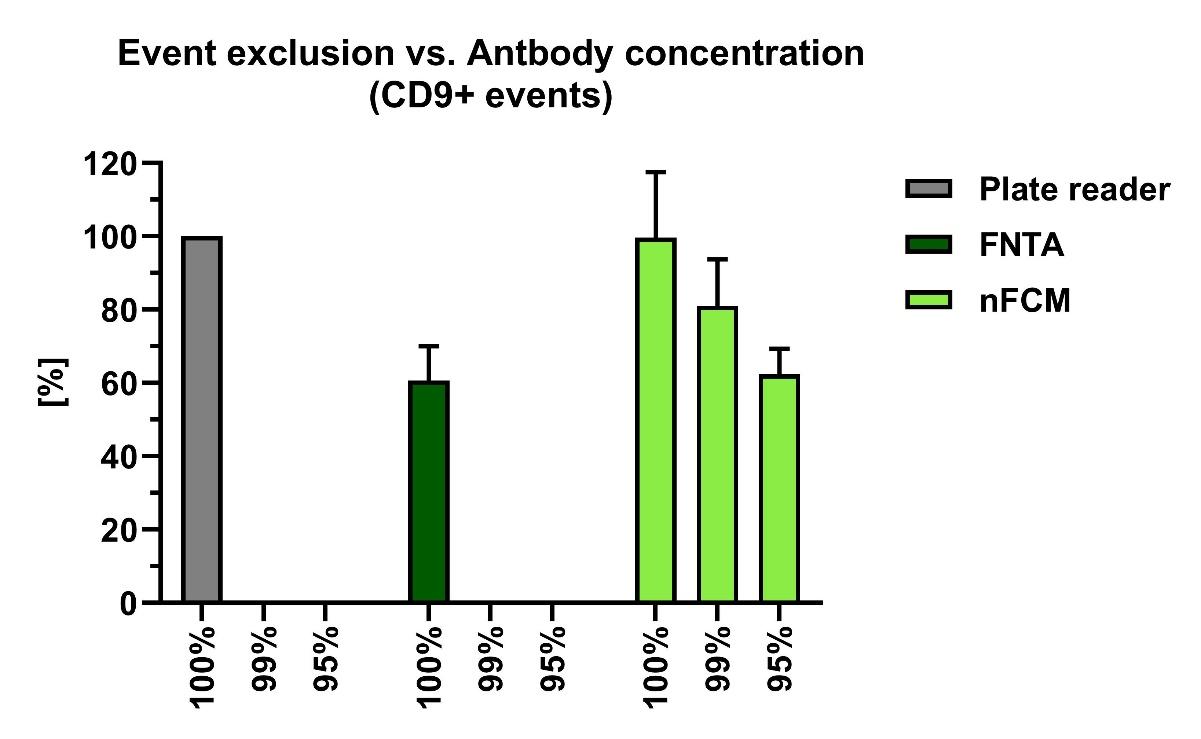

**Figure S4. Exclusion of the brightest events significantly impacts antibody concentration measurements.** nFCM gating strategy led to inclusion of fewer brightest events which significantly contributed to the estimation of antibody concentration and made it more comparable to the bulk measurements. Provided that the gating strategy excluded 1% or 5% of the top brightest events, this would have resulted in an exclusion of 18% or 37% of antibodies from the analyses, respectively, and made the latter more comparable to the FNTA results. This showcases the extremely uneven (non-Gaussian) distribution of epitopes across the whole COLO EV population.

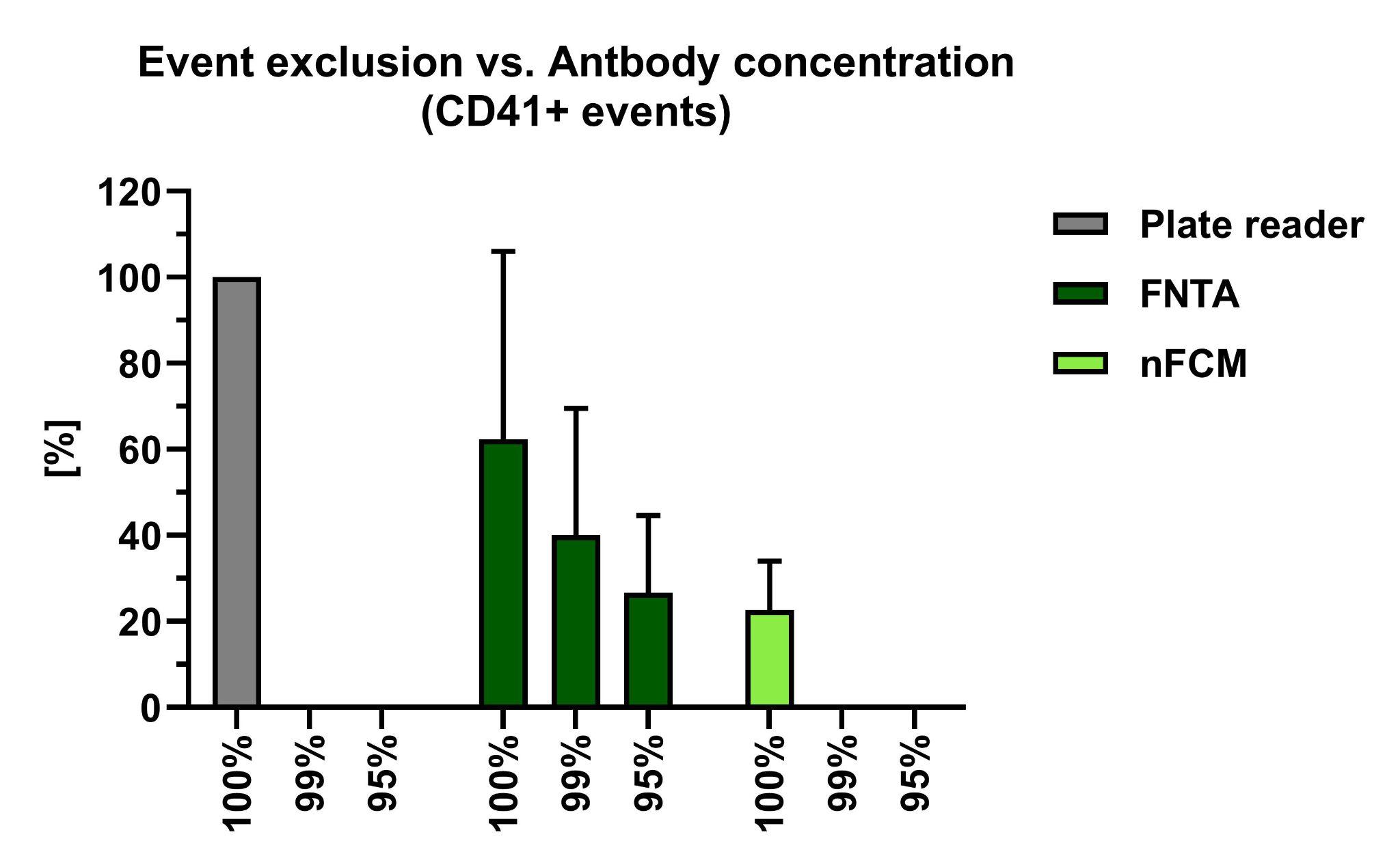

**Figure S5. Exclusion of brightest events significantly impacts antibody concentration measurements.** FNTA’s inability to detect and track dimmer events led to biased analyses of fewer brightest events which significantly contributed to the estimation of antibody concentration and made it more comparable to the bulk measurements. Provided that the gating strategy excluded 1% or 5% of the top brightest events, this would have resulted in an exclusion of 22% or 36% of antibodies from the analyses, respectively, and made the latter more comparable to the nFCM results. This showcases the extremely uneven (non-Gaussian) distribution of epitopes across the whole plasma EV population.

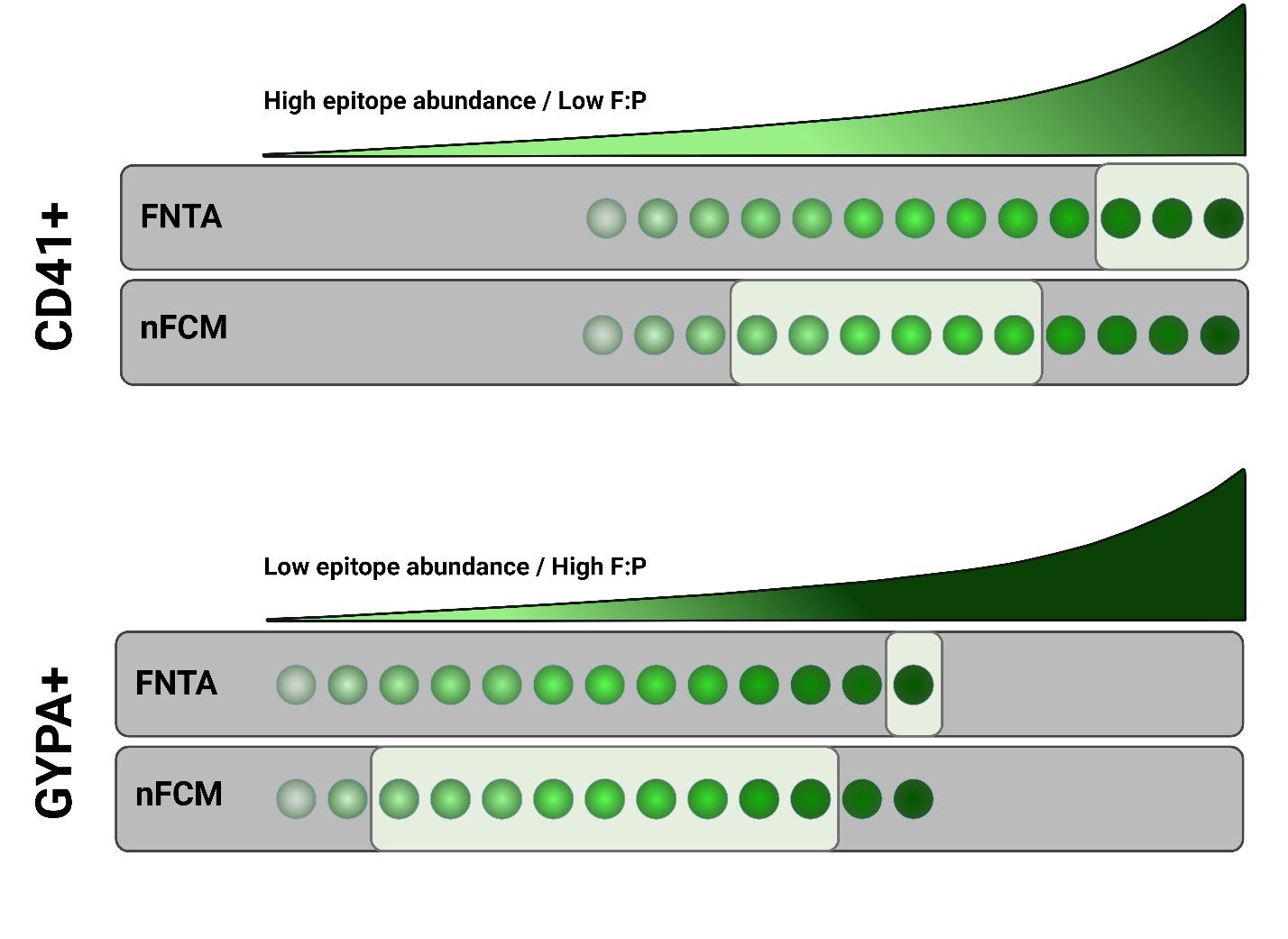

**Figure S6. Multiple factors affect the detection of the fluorescent events.** Epitope abundance, F:P ratio of the antibody used, gating strategy, and instrument’s sensitivity have a combined effect on the detectability of fluorescent events. The exacerbated schematic model illustrates this effect, taking as an example CD41+ and GYPA+ plasma EVs analyzed on FNTA and nFCM. Note that the CD41+ EV with more epitopes might appear as bright as GYPA+ EV with less epitopes, due to the different F:P ratios of these antibodies. Anti-CD41 antibody had <3 conjugated AF488 molecules, while anti-GYPA antibody had ~5 conjugated AF488 molecules, making it almost twice as bright. Created with Biorender.
